# Gene function contributes to gene expression levels in *S. cerevisiae*

**DOI:** 10.1101/330365

**Authors:** Mark J. Hickman, Andrea Jackson, Abigail Smith, Julianne Thornton, Amanda Tursi

## Abstract

It is not understood what evolutionary factors drive some genes to be expressed at a higher level than others. Here, we hypothesized that a gene’s function plays an important role in setting expression level. First, we established that each *S. cerevisiae* gene is maintained at a specific expression level by analyzing RNA-seq data from multiple studies. Next, we found that mRNA and protein levels were maintained for the orthologous genes in *S. pombe*, showing that gene function, conserved in orthologs, is important in setting expression level. To further explore the role of gene function in setting expression level, we analyzed mRNA and protein levels of *S. cerevisiae* genes within gene ontology (GO) categories. The GO framework systematically defines gene function based on experimental evidence. We found that several GO categories contain genes with statistically significant expression extremes; for example, genes involved in translation or energy production are highly expressed while genes involved in chromosomal activities, such as replication and transcription, are weakly expressed. Finally, we were able to predict expression levels using GO information alone. We created and optimized a linear equation that predicted a gene’s expression based on the gene’s membership in 161 GO categories. The greater number of GO categories with which a gene is associated, the more accurately expression could be predicted. Taken together, our analysis systematically demonstrates that gene function is an important determinant of expression level.

## INTRODUCTION

Proteins play critical roles in cellular metabolism, structure and homeostasis. Each step of gene expression is intricately regulated to ensure that the abundance of each protein is appropriate for the cellular condition (Wittkopp 2014). With recent advances in protein quantitation, it has been possible to ascertain “absolute” protein abundances (Vogel and Marcotte 2012; Liu *et al*. 2016). These technologies have revealed that the steady-state abundance of each protein remains similar across studies (Conesa *et al*. 2016), suggesting that there is a set point for each protein. The steady-state abundance of each protein is highly correlated with that of orthologous proteins across diverse taxa (Schrimpf *et al*. 2009; Laurent *et al*. 2010; Khan *et al*. 2013). Since orthologs are known to share function (Dolinski and Botstein 2007), the fact that protein abundance is widely conserved suggests that the function of each protein is important in determining its abundance.

It is expected that, through evolutionary forces, protein abundance reaches a level that maximizes the fitness of the organism. Two opposing factors influence the expressed level of a protein: the cost of protein synthesis drives down expression while the biochemical need for the protein drives up expression, ultimately resulting in a level that maximizes fitness (Wagner 2005; Dekel and Alon 2005; Lang *et al*. 2009). We hypothesize that this biochemical need can be predicted by the protein’s function, as captured in the gene ontology (GO) framework. Taking this one step further, we propose that the GO terms describing a gene product can be used to predict protein abundance. Thus, genes that share a GO term will exhibit similar protein abundances because the proteins work with related biochemical parameters. Gene ontology (GO) attempts to define three aspects of a gene’s function (molecular function, biological process, and cellular component) and these aspects can be used to think about how function may influence expression level. For example, proteins with the same molecular function (e.g. isomerase activity) may have comparable Michaelis-Menten kinetics (e.g., K_m_, k_cat_) and work on substrates with related concentrations. Proteins that participate in the same biological process (e.g., cytoplasmic translation) are components of a pathway that may have similar flux at each step. Proteins within the same cellular component (e.g., nucleus) are confined to the same physical volume. Supporting the idea that gene function determines abundance, it has been shown in genome-wide mRNA and protein studies that transcription factors exhibit low abundance (Drawid *et al*. 2000; Ghaemmaghami *et al*. 2003; Vaquerizas *et al*. 2009) while protein synthesis and metabolism genes exhibit high abundance (Velculescu *et al*. 1997; Jansen and Gerstein 2000; Nagalakshmi *et al*. 2008).

In this study, we took a systematic genome-wide approach to investigate whether *S. cerevisiae* gene function (as indicated by gene membership in GO categories) is related to gene expression level. *S. cerevisiae* is suitable for this study because cell type-specific expression is not an issue, several genomic studies of RNA and protein abundance have been performed, and a large proportion of genes have been well-annotated. As an indicator of expression level, we mainly relied on mRNA abundance (though we confirmed some of our findings with protein abundance data) because protein abundance measurements are not consistent between studies and are limited in their genome-coverage (Vogel 2013; Liu *et al*. 2016). Moreover, with the recent improvement of data quality, it has been found that mRNA levels are strong predictors of protein levels (Csárdi *et al*. 2015; Li *et al*. 2017), in contrast to earlier studies showing a weaker correlation between mRNA and protein abundance (Maier *et al*. 2009). In this study, we found that mRNA and protein levels of *S. cerevisiae* genes are highly correlated to levels of orthologous genes in *S. pombe*, supporting the notion that gene function, which is shared among orthologues, determines expression level. Then, we statistically analyzed the set of genes within each of 161 GO categories and found that many GO categories exhibit statistically significant expression extremes. For example, genes involved in translation or the cell wall are highly expressed while genes involved in chromosomal activities, such as replication and transcription, are weakly expressed. Furthermore, we wanted to test whether GO categories could be used to predict gene expression so we developed and optimized a linear model in which GO categories could be used to determine expression. Using this method, we were able to predict expression of *S. cerevisiae* and *S. pombe* genes with GO category information alone. Together, these data show that the function of a gene is a determinant of its expression level, adding to our understanding of the evolution of gene expression.

## MATERIALS AND METHODS

### *S. cerevisiae* datasets

RNA-seq datasets were downloaded from NCBI Gene Expression Omnibus (GEO) or Sequence Read Archive (SRA). *S. cerevisiae* sets included SRA048710 (Risso *et al*. 2011), GSE43002 (Baker *et al*. 2013), GSE61783 (Adhikari and Cullen 2014), GSE52086 (Martín *et al*. 2014), GSE57155 (Fox *et al*. 2015), and GSE85595 (Bendjilali *et al*. 2017). Datasets that were published as SRA files were converted to FASTQ files with the SRA toolkit (Leinonen *et al*. 2011), trimmed with the FASTQ Quality Trimmer (Blankenberg *et al*. 2010) using a quality score of ten, mapped to the R64.1.1 2011-02-03 yeast genome (Engel *et al*. 2014) with TopHat2 (Kim *et al*. 2013), and converted into raw counts per gene with HTSeq (Anders *et al*. 2014). Gene counts were normalized for gene length and the total number of sequencing reads, thus generating RPKM (Reads Per Kilobase of transcript per Million mapped reads) (Mortazavi *et al*. 2008). In studies that did not provide the total number of mapped reads, the total number of reads that mapped to genes was used. Each replicate within a study was treated individually when averaging all replicates together; there were 18 total *S. cerevisiae* RNA-seq replicates. Protein abundance in *S. cerevisiae* was determined by mass spectrometry (Lawless *et al*. 2016). Paralogous proteins could not be distinguished in this study so were removed from our analysis, leaving absolute abundances for 1103 unique proteins. Gene names in all datasets were converted to systematic gene names using gene names from the Saccharomyces Genome Database (SGD) (Cherry *et al*. 2012). Cell cycle data generated in a separate study (Spellman *et al*. 1998) were downloaded from SGD. File S1 contains *S. cerevisiae* gene names and normalized expression data. File S5 contains gene ontology (GO) SLIM categories for each gene, downloaded from SGD.

### *S. pombe* expression datasets

*S. pombe* gene names and their respective *S. cerevisiae* orthologues were obtained from PomBase (Wood *et al*. 2012; McDowall *et al*. 2015). *S. pombe* RPKM values were averaged from two RNA-seq datasets: GSE74411 (Mukherjee *et al*. 2016) and GSE80349 (Shah *et al*. 2016). Protein abundance in *S. pombe* was determined by integrating data from several studies (Wang *et al*. 2015); abundance data were downloaded from the Protein Abundance Database (PaxDb). File S2 contains *S. pombe* gene names and normalized expression data.

### Expression of each GO category

The expression of genes within each GO category (e.g., “mRNA processing”) was summarized by four metrics: mean RPKM, median RPKM, mean rank median rank. For ranks, GO categories were ordered from least to highest RPKM (mean or median) and given a rank. Using these metrics, the expression of each GO category was compared to that of all genes. To determine whether the GO category differed significantly from all genes, the metric was compared to the respective metric of a randomly-selected set of the same size. This comparison was performed with 10^7^ iterations, counting the number of iterations that resulted in a metric more extreme than the original metric for the category. More extreme refers to either tail of the distribution, depending on whether the original metric for the category was higher or lower than all genes. Thus, *p* = (number of iterations resulting in a more-extreme metric) / 10^7^, and the p-value refers to the probability that the expression of genes within a GO category is either higher or lower than all genes by chance. This was performed in the R statistical language (Team 2015), as shown in File S6. The p-values were corrected for multiple hypothesis testing, using the BH method (Benjamini and Hochberg 1995).

### Prediction

The RPKM level of each gene was predicted based on its inclusion in each of the 163 GO categories, according to this linear equation:

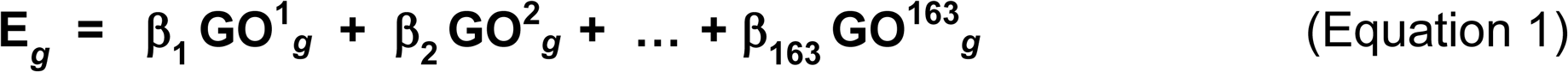

where E_*g*_ represents the predicted expression level for gene, *g*; β_1_ through β_163_ are 163 coefficients to be optimized; and GO^1^_*g*_ through GO^163^_*g*_ are binary numbers signifying whether gene *g* is present (=1) or absent (=0) in the respective GO category.

The 163 β values were adjusted over 10^6^ iterations using a random walk, with the goal of maximizing the correlation between the predicted expression (log_10_) and the actual RPKM (log_10_) for all genes. To begin, one of the 163 β values was randomly chosen and then changed randomly up or down, with a step size of 1. If the change increased the correlation, then this specific change was repeated. To avoid reaching a local maximum, the change was repeated only 90% of the time. If the change decreased the correlation, then another random β value was chosen and changed. The iterations continued until a maximum correlation was achieved. This was carried out in R, as shown in File S7.

### Data availability

File S1 lists *S. cerevisiae* genes and associated expression and cell-cycle data. File S2 lists *S. pombe* genes and associated orthologue and expression data. File S3 lists GO categories and associated expression and statistics data. File S4 shows starting seeds and resulting beta values for 10 independent random walks. File S5 lists *S. cerevisiae* genes and their GO Slim categories, adopted from www.yeastgenome.org. File S6 is the R script which determines the significance of gene expression within each GO category. File S7 is the R script which performs a random walk to predict expression. Figure S1 depicts mRNA abundance across studies. Figures S2, S3, and S4 show the expression of all genes within 100 GO processes, 40 GO molecular functions, and 21 GO cellular components, respectively. Figure S5 depicts the expression of cell cycle genes vs. non-cell cycle genes. Figure S6 shows the expression of genes within specific cell-cycle phases. Figure S7 shows that RPKM is correlated to protein abundance.

Figure S8 compares the protein and RPKM datasets, regarding the number of genes per GO category. Figure S9 shows two independent random walks, to predict RPKM levels, generated similar β coefficients. Figure S10 graphs predicted vs. actual expression, with the “Cytoplasmic translation” genes colored in red. Figure S11 graphs predicted vs. actual expression of “Cytoplasmic translation” genes, with genes exhibiting an identical predicted expression value of 197.8 colored in red. Figure S12 shows the distribution of how many GO categories describe each gene. Figure S13 shows that GO categories can be used to predict protein abundance, using a random walk. Figure S14 shows two independent random walks, to predict protein abundance, generated similar β coefficients. These files have been submitted to https://gsajournals.figshare.com/submit.

## RESULTS

### *S. cerevisiae* genes exhibit an expression set point

We first evaluated whether each *S. cerevisiae* gene exhibits consistent steady-state mRNA abundance relative to all other genes. We monitored mRNA levels across eighteen independent samples drawn from six RNA-seq studies using the standard lab strains, S288C and Sigma, grown in rich media at 30°C. Transcript levels were calculated as RPKM values (Reads Per Kilobase of transcript per Million mapped reads), which have the benefit of allowing between-gene comparisons of mRNA abundance (Mortazavi *et al*. 2008). Figure S1 shows that the mRNA abundance of each gene is highly correlated across studies (r = 0.69 to 0.93). These results indicate that each gene is maintained at an expression “set point.”

### *S.cerevisiae* expression levels are correlated with orthologous genes in *S. pombe*

It has been shown that orthologous genes, which share function, between a diverse set of species including *S. cerevisiae*, *C. elegans*, *D. melanogaster* and *H. sapiens* exhibit highly similar abundances of both RNA and protein (Schrimpf *et al*. 2009; Laurent *et al*. 2010; Khan *et al*. 2013), supporting the idea that function has an influence on abundance. We confirmed that this is the case even when comparing orthologous genes between *S. cerevisiae* and *S. pombe*, two species separated by 330-420 million years of evolution (Sipiczki 2000). Both mRNA and protein levels of orthologous genes are highly correlated between these yeast species (Figure 1). This result further supports the hypothesis that gene function, which is conserved among orthologues, is important in determining expression levels.

**Figure 1.**
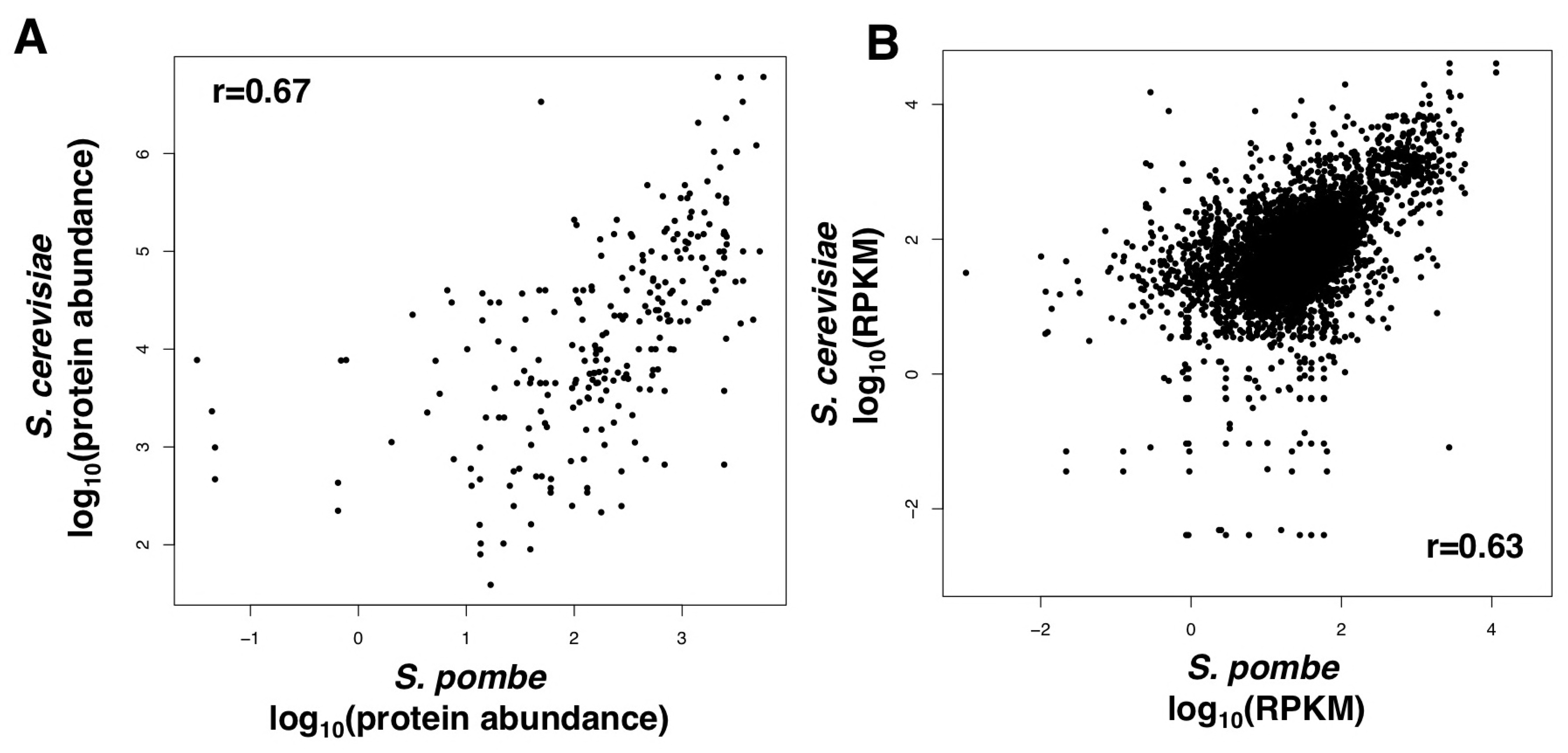
The expression of orthologous genes in *S. cerevisiae* and *S. pombe* are highly correlated. **(A)** Comparison of protein abundances between orthologous genes in *S. pombe* and *S. cerevisiae* (r=0.67, n=111). **(B)** Comparison of RPKM values between orthologous genes in *S. pombe* and *S. cerevisiae* (r=0.63, n=1118).

### Evaluating the expression level within each GO category

To more deeply explore how gene expression levels are related to gene function, we employed the GO Slim annotations at the Saccharomyces Genome Database (SGD) (Cherry *et al*. 2012). In the GO framework, experimental evidence is used to associate each gene with one or more GO annotations which describe biological processes, molecular functions, and cellular components. The SLIM annotations used here are unique to *S. cerevisiae* and were developed by SGD to broadly categorize genes into their functional groups. We employed this condensed set of annotations in order to test whether expression level varies across these broad functional categories.

We characterized the distribution of mRNA expression levels of genes associated with 100 biological processes (Figure 2A), 40 molecular functions (Figure 2B), and 21 cellular components (Figure 2C). Organizing the categories into three panels facilitates appropriate comparisons; for example, comparing the two components “nucleus” and “cell wall” is more appropriate than comparing the component “nucleus” to the process “mRNA processing.” Each panel of Figure 2 is ordered by the median expression level of a GO category. Also shown are the mean expression levels, to test whether there is a skewed distribution of expression within categories. The complete distribution of expression levels within each GO category is depicted in Figures S2, S3 and S4. These figures show that, while there is wide variation of expression values within each GO category, some GO categories exhibit higher expression levels than others.

**Figure 2.**
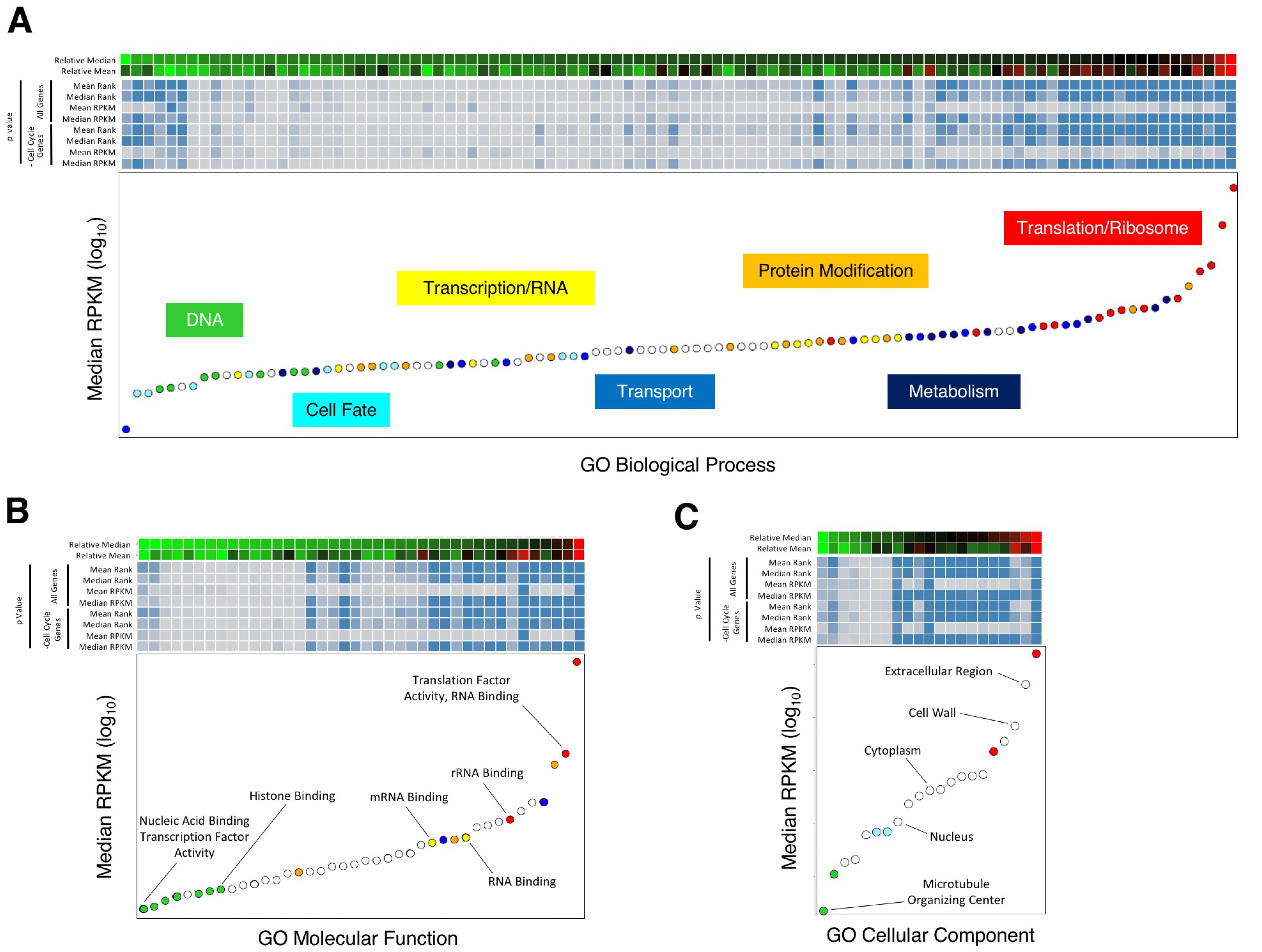
The mRNA abundance of genes within each GO Category: **(A)** Biological Processes, **(B)** Molecular Functions, and **(C)** Cellular Components. Each panel is organized in the same manner. The GO categories are ordered from left-to-right with increasing RPKM median. The first two rows are in heatmap format and depict the mean and median, respectively, of each GO category; black signifies genome-wide mean or median, red signifies higher than genome, and green signifies lower than genome. Rows 3-6 are in heatmap format and depict the probabilities that the mean or median differs from the genome-wide mean or median by chance, using random sampling with four different metrics; as the intensity of blue increases, the p-value decreases (minimum *p*=10^-7^). Removing the 799 cell cycle genes from the analysis had little effect on the probabilities (rows 7-10). Finally, the median RPKM (log_10_) of each GO category was plotted; GO categories representing related functions (e.g., DNA) were merged into groups and colored, as described in the Materials and Methods and in File S3.

To test whether the distribution of expression levels in each GO category is significantly lower or higher than expected, we compared expression of genes within each GO category to a set randomly selected from the genome, as described in the Materials and Methods. This one-sided test generated a p-value for each GO category, equal to the probability that the distribution of expression values could occur by chance. The results are shown in heatmap format in Figure 2, with darker blue indicating greater significance (i.e., lower p-value). To ensure that the statistics are robust, four metrics were used to describe the distribution of expression levels in each GO category: median RPKM, mean RPKM, median rank, and mean rank. Three of the metrics (median RPKM, median rank, and mean rank) resulted in p-values that are remarkably similar, as shown by a similar shading of blue in Figure 2. In contrast, the mean RPKM metric often resulted in higher p-values. A likely explanation is that the mean RPKM of each category is influenced by outlier expression values, making the comparison to all genes less meaningful. It was for this reason that we decided to employ median and rank, in addition to mean, as metrics in our analyses.

We were concerned that our analyses would be biased by cell-cycle genes, which may have low expression because their expression is limited to a subset of the cell cycle. Surprisingly, the cell cycle genes, as identified previously (Spellman *et al*. 1998), do not appear to be expressed less than non-cell-cycle genes (Figure S5). Upon closer examination, we found that the G_1_ genes are expressed significantly less than non-G_1_ genes (Figure S6). To rule out any cell-cycle effects on gene expression within GO categories, we repeated the statistical tests on the entire gene set minus the 799 cell-cycle genes (Figure 2). The p-values obtained were largely unchanged, as shown by the similar shading of blue in the heatmap, indicating that our results are not biased by cell-cycle genes.

Now that we have described the methodology employed to analyze expression within each GO Category, we will use the next three sections to explore how expression levels relate to (1) Biological Processes, (2) Molecular Functions, and (3) Cellular Components.

### Expression of genes within GO Biological Processes

Figure 2A depicts the expression of the 100 GO Biological Processes ordered by median RPKM levels. Also, see File S3 for category statistics and Figure S2 to visualize the distribution of gene expression within each category. Several notable patterns are observed when considering the expression level within GO categories. To facilitate description of these patterns, we have grouped related GO terms (as shown by the colored points in Figure 2). First, there is a clear relationship between the GO terms and the Central Dogma (i.e., DNA → RNA → protein). Among the lowest expressed GO terms are those involving DNA processes (indicated by green points), such as “chromosome segregation” and “DNA repair.” Our statistical tests show that these categories exhibit significantly low expression. This is followed by terms describing aspects of transcription and RNA processes, such as “mRNA processing” and “transcription from RNA polymerase II promoter” (indicated by yellow points). The GO terms showing statistically high expression are related to aspects of translation and the ribosome (indicated by red points). For this group, we included certain transcription terms (e.g., “transcription from RNA polymerase I promoter”) because these processes solely serve to create structural RNAs of the ribosome. The relationship between gene expression level and the role of the gene in the Central Dogma can be explained by “amplification”. In *S. cerevisiae*, experimental data show that when mRNA was detected for ∼5854 genes, there were ∼36,000 total mRNA molecules and 35 million proteins per haploid cell (Csárdi *et al*. 2015). Thus, the amplification from DNA to mRNA is 6-fold while the amplification from mRNA to protein is 972-fold. Another study measured the components of cell dry weight, which includes abundant rRNA molecules, and found that DNA amount is 20-fold less than RNA amount and that RNA amount is 5-fold less than protein amount (Feijó Delgado *et al*. 2013). As might be expected, our findings suggest that the measured cellular concentration of these biomolecules (i.e., DNA, mRNA, protein) is related to the expression level of the proteins that are tasked with synthesizing or maintaining the respective biomolecule.

The genes that participate in protein modification (orange points; e.g., “protein modification” and “protein acylation”) exhibit relatively high expression, but less so than the translation and ribosome genes. There could be two reasons for this, not necessarily mutually exclusive. First, each modification enzyme may only work on a subset of proteins while translation/ribosome proteins work on all proteins. Second, modification enzymes catalyze only one or few reactions on each polypeptide substrate while each translation/ribosome protein contributes to dozens or hundreds of peptide bond formation reactions to create a single polypeptide. In both cases, a modification enzyme is likely less expressed than a translation/ribosome protein due to decreased flux through the enzyme.

Genes involved in several GO metabolic processes (dark blue points), such as “nucleobase-containing small molecule metabolic process,” are highly expressed, consistent with the large flux occurring through biosynthesis and energy production pathways. Interestingly, some metabolic processes, like “oligosaccharide metabolic process,” contain genes of low expression. This makes sense because the mRNA expression levels observed here are from yeast grown in the monosaccharide glucose as the carbon source, not from yeast grown in oligosaccharides.

The GO categories associated with transport (blue points) deserve mention because they are among the highest and lowest expressed. The highest among the Transport categories is “nucleobase-containing compound transport”, which facilitates the much-needed transport of nucleobases for metabolism. The next highest category is “nuclear transport” comprised of genes involved in the nuclear pore and in export of ribosomal RNA, both important for translation. On the other hand, certain Transport categories, such as “carbohydrate transport,” exhibit low expression, which is not surprising given that the cells were grown in excess glucose, which represses certain types of sugar import (Ozcan and Johnston 1999).

Finally, GO categories related to Cell Fate (light blue points) mainly show low expression. This is consistent with the cells being cultured under asexual rich-media conditions and thus not faced with cell fate decisions (e.g., meiosis, mating, invasive growth). Surprisingly, “mitotic cell cycle” exhibited low expression, despite the cells undergoing exponential growth. A possible factor is that many of the cell cycle genes are involved in signal transduction and transcriptional regulation, both of which exhibit low expression. Specifically, “signaling” is 36^th^ lowest out of 100 GO Processes, and “nucleic acid binding transcription factor activity” is the lowest out of 40 GO Functions.

### Expression of genes within GO Molecular Functions

Figure 2B shows the expression of the 40 GO Molecular Functions ordered by median RPKM levels. Also, see File S3 for category statistics and Figure S3 to visualize the distribution of gene expression within each category. Again, the relationship between GO terms and the Central Dogma is apparent. Among the lowest expressed GO terms are those involving DNA processes (green points), such as “nuclease activity” and “DNA binding.” We included “nucleic acid binding transcription factor activity” and “transcription factor binding” in the DNA group because the proteins (transcription factors and their regulators) within these categories mainly bind to DNA. The proteins do not participate in the high-flux enzymatic steps of transcription and RNA processing, but instead bind to a limited number of sites on the DNA. Generally exhibiting higher expression are categories such as “RNA binding” that describe aspects of transcription and RNA processes (yellow points). Finally, GO terms such as “translation factor activity, RNA binding” that describe aspects of translation and the ribosome exhibit the highest expression (red points).

Like the GO processes in Figure 2A, the GO Functions that are involved in protein modification (indicated by orange points in Figure 2B) exhibit relatively high expression but reduced expression compared to translation and ribosome categories. An example is “unfolded protein binding,” the third highest Function, which can be compared to the related “protein folding,” the fifth highest Process.

Also notable are the two transport-related GO Functions (“protein transporter activity” and “transmembrane transporter activity”) which are relatively highly expressed (blue points). There are only two transport-related Functions compared to 10 transport-related Processes (Figure 2A). When observing the 10 Processes, there is much more variation in expression levels, indicating that some of this variation is lost when grouping genes into only 2 categories.

### Expression of genes within GO Cellular Components

Figure 2C shows the expression of the 21 GO Cellular Components ordered by median RPKM levels. Also, see File S3 for category statistics and Figure S4 to visualize the distribution of gene expression within each category. Consistent with our previously established relationship between expression and the Central Dogma, categories involving DNA processes (“microtubule organizing center” and “chromosome,”) show low expression while categories involving Translation/Ribosome (“nucleolus” and “ribosome,”) exhibit high expression. There are no Cellular Component categories that capture only RNA/Transcription genes.

Categories related to Cell Fate (“cellular bud” and “site of polarized growth,”) exhibited relatively low expression, possibly because these cellular locations are short-lived and comprise a small space. As expected, the genes within the cytoplasm are expressed at higher levels than genes within the nucleus, a cellular component with a volume substantially less than that of the cytoplasm (Jorgensen *et al*. 2007).

Additionally, genes within the categories “Extracellular Region” and “Cell Wall” are highly expressed, likely due to the vast number of proteins needed to populate these spaces (de Groot *et al*. 2009).

### Gene function is also associated with protein expression

So far, we have shown that mRNA levels are correlated with gene function. Since proteins actually carry out the function, we also attempted to associate protein levels with function. This task was hindered because quantitative data for protein levels is lacking. Not only is it difficult to detect levels for many proteins, abundance measurements are not consistent between the limited number of studies (Vogel and Marcotte 2012; Liu *et al*. 2016). We identified one recent study that determine the absolute abundances of 1103 *S. cerevisiae* proteins with high-quality by using mass spectrometry with internal controls (Lawless *et al*. 2016). We found that the measured protein abundance is highly correlated (r=0.61) with the RPKM values that we used here (Figure S7). Next, we compared protein expression within each GO category with mRNA expression within each category (Figure 3A). There was only a modest correlation (r=0.44). We hypothesized that the low correlation is due to the small number of genes in the protein dataset and the resulting smaller number of genes per GO category (Figure S8). To control for this discrepancy in number of genes per category, we calculated the median RPKM for each GO category using only the 1103 genes that are in the protein dataset. Then, we compared the RPKM of each GO category with the corresponding protein abundance (Figure 3B). There was a high correlation (r=0.81), suggesting that gene function, as defined by GO categories, has an effect not only on mRNA levels but on protein levels as well.

**Figure 3.**
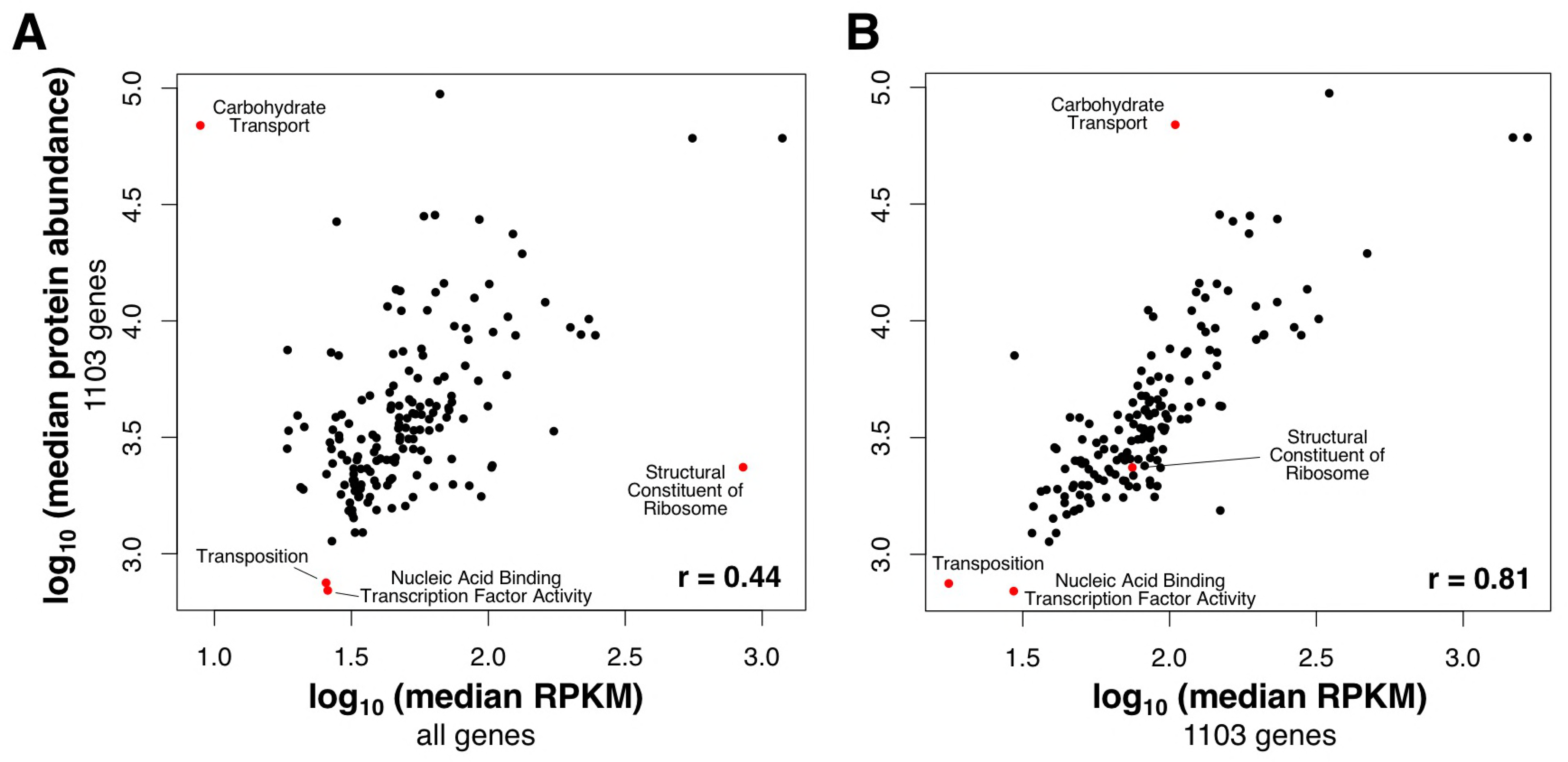
The median mRNA abundance of each GO category is similar to the median protein abundance. **(A)** For each GO category, the median protein abundance is plotted against the median RPKM. Note that the protein abundance dataset only includes 1103 genes. **(B)** For each GO category, the median RPKM abundance (calculated from the 1103 genes in the protein data set) is plotted against the median protein abundance.

### Gene function can predict expression levels

As described above, we found that the genes in each GO Category have distinct expression levels. We wondered whether gene function, as assessed by a gene’s membership in GO categories, can be used to predict expression level. To test this, we developed a linear equation in which the RPKM of each gene is determined by the gene’s inclusion in each of the 163 GO categories (see Materials and Methods). Each GO category was assigned a coefficient (β), which was optimized using a random walk, with the goal of accurately predicting the expression of each gene. Prediction accuracy was assessed by correlating the predicted vs. the actual expression of all genes. As shown in Figure 4A, as the random walk progressed over 10,000 iterations, the correlation increased until a maximum of 0.44 was reached. The correlation did not increase further, even when the walk was performed with 10^7^ iterations (data not shown). Additionally, when the random walk was initiated 10 independent times with randomly-chosen β coefficients, the same correlation (0.44) and β coefficients were obtained (File S4). For example, Figure S9 shows a linear relationship between the β coefficients of the 2^nd^ and 3^rd^ repeats. As might be expected, the GO categories that had a high coefficient (e.g., “cytoplasmic translation”) were among the highest expressed categories. In contrast, the GO categories that had a low coefficient (e.g., “cellular respiration”) were not always the lowest expressed categories; this suggests that these categories, in combination with other GO categories, play a complicated and additive role in prediction.

**Figure 4.**
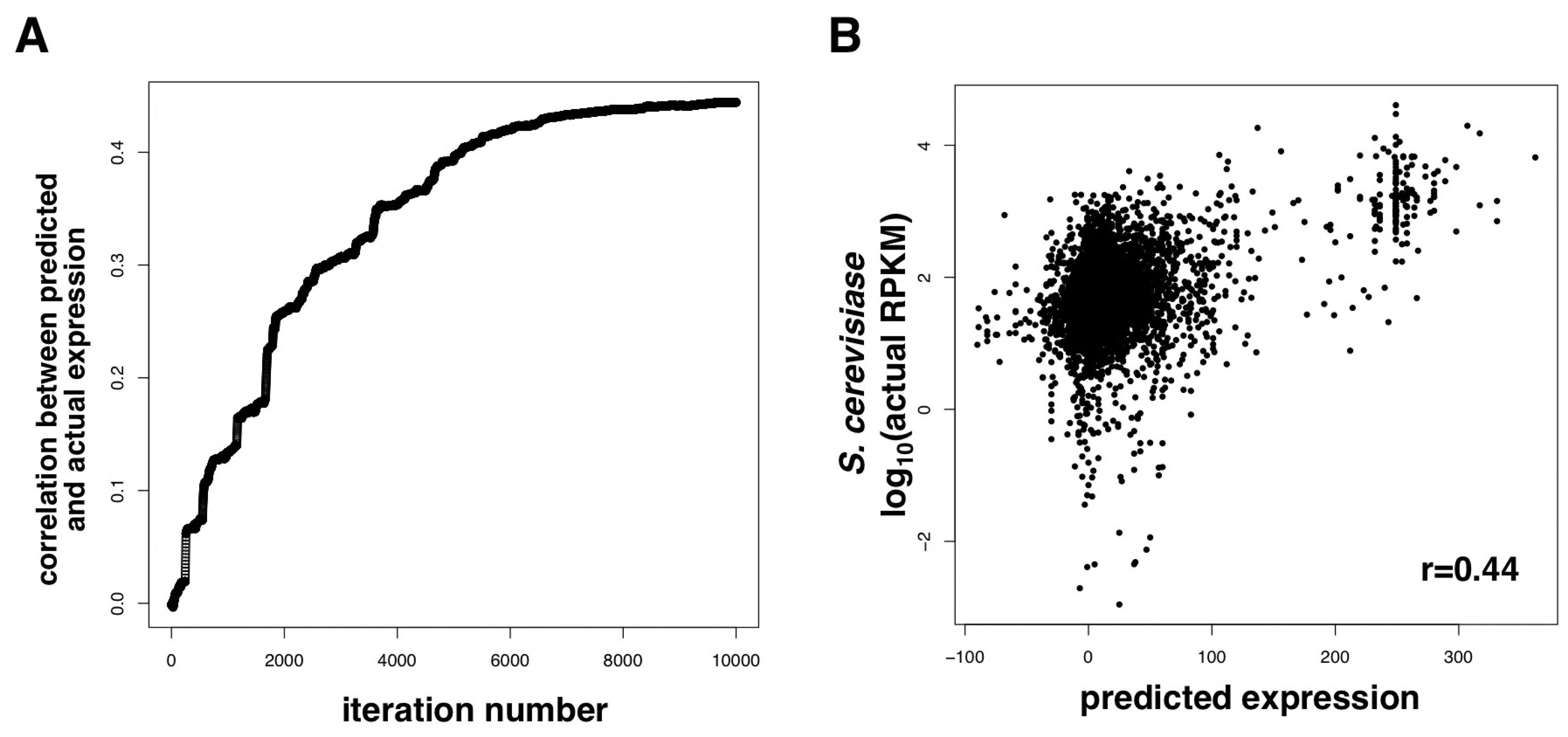
GO categories alone can be used to predict gene expression levels. **(A)** A random walk was performed to optimize the β coefficients of each GO category in predicting expression, as described in the Materials and Methods. The graph depicts the iteration number vs. the correlation between predicted and actual RPKM values of all genes. **(B)** Shown is the predicted expression (x-axis with arbitrary scale) vs. the actual log_10_(RPKM) in *S. cerevisiae*.

The predicted expression vs. actual expression of each gene is depicted in Figure 4B. This graph shows that there is a modest correlation between predicted and actual expression. A notable feature of the graph is that some genes can be grouped together into a vertical line; in such a group, each gene has the same predicted expression but a variety of actual expression levels. This is likely caused by having incomplete functional information; the genes are predicted to have the same expression level because they are in the same GO category and are not functionally differentiated by other informative GO categories. An interesting example is the set of genes in the “cytoplasmic translation” category (Figure S10). As expected, these genes are predicted to have high expression. A subset of these genes was predicted to have identical expression but actually vary in expression (Figure S11). The reason that these genes are predicted to have the same expression level is that they share membership in the same 5 GO categories (cytoplasm, cytoplasmic translation, ribosome, structural constituent of ribosome, structural molecule activity). If these genes had additional functional information, the prediction would likely be more accurate.

To further test this idea, we limited our prediction to genes associated with a minimum number of GO categories. Genes show a wide-range in the number of GO categories assigned to them (Figure S12), from 0 to 35, presumably attributable to the degree to which the genes have been studied. As we expected, when the prediction was limited to genes associated with a larger number of GO categories, the prediction increased in accuracy, as shown by a higher correlation between predicted and actual expression (Figure 5). It should be noted that as the minimum number of GO categories increases, the number of genes dramatically decreases (blue line in Figure 5). Regardless, these results suggest that having more information about gene function improves the ability to predict gene expression levels.

**Figure 5.**
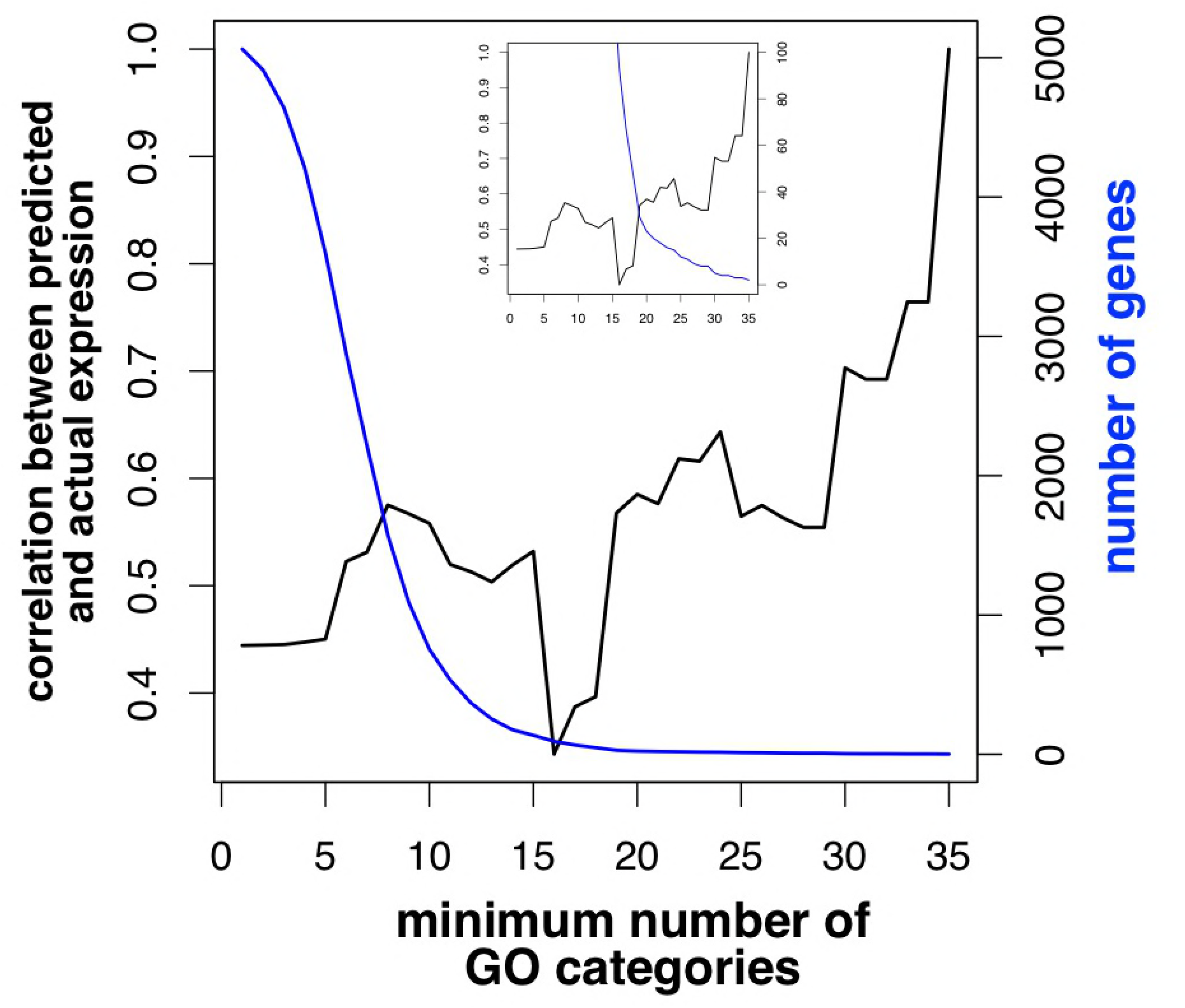
Limiting prediction to genes with a minimum number of GO categories improves the correlation between predicted and actual expression (black line). Also, as the minimum increases, the number of genes meeting or surpassing this minimum decreases (blue line). In the inset graph, the range of the “number of genes” axis is 0 to 100.

In our prediction analysis above, we created a model that predicts gene expression based on gene function. We wanted to test whether this model, developed with *S. cerevisiae* GO annotations, can be used to predict expression levels of the orthologous genes in *S. pombe*. Indeed, we found that there was a high correlation (r=0.54) between our predicted expression values and actual expression in *S. pombe* (Figure 6).

**Figure 6.**
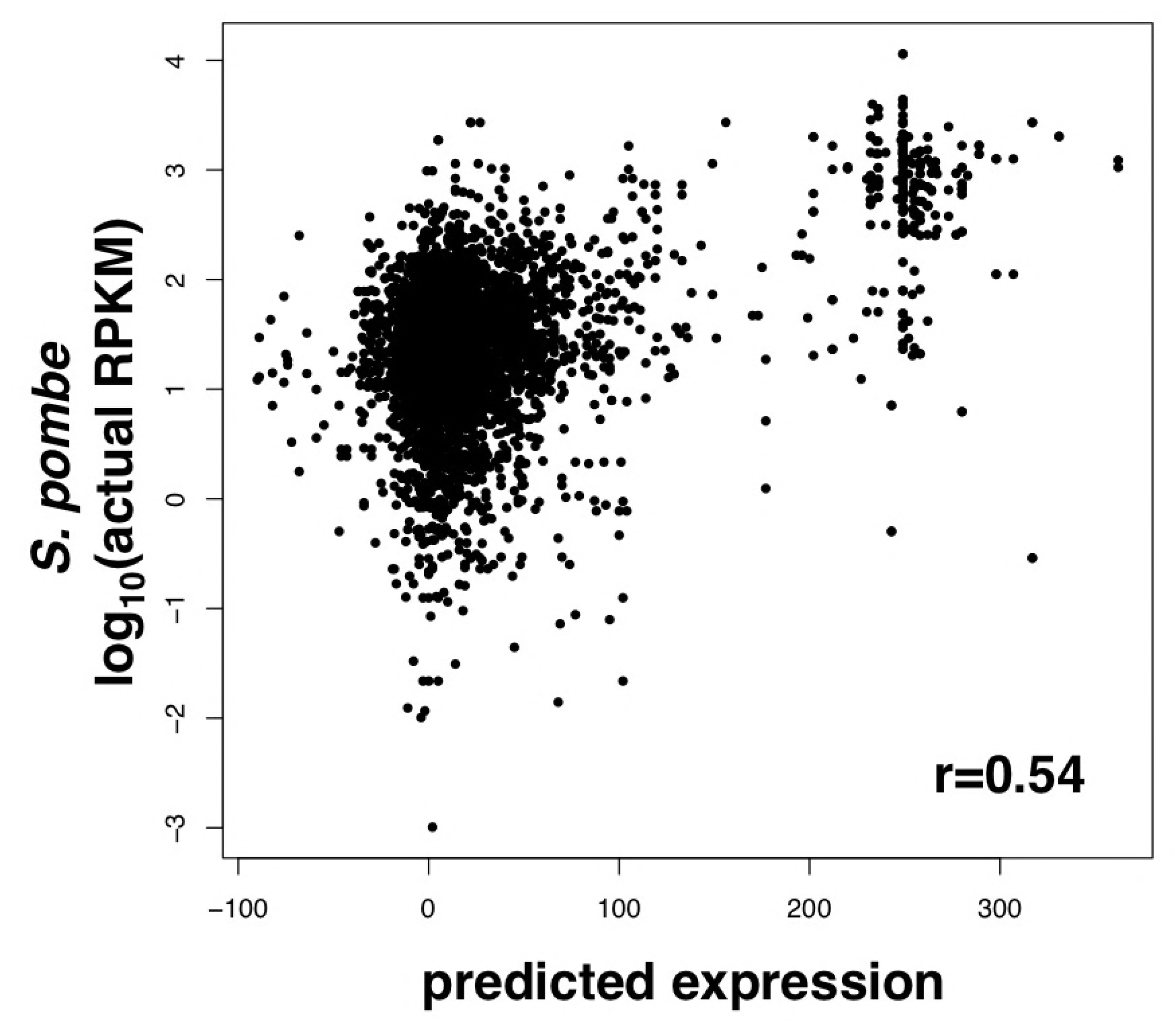
Gene function in *S. cerevisiae* can be used to predict expression of orthologues genes in *S. pombe*. Shown is the predicted expression of *S. cerevisiae* genes (x-axis with arbitrary scale) vs. the actual log_10_(RPKM) of their respective orthologs in *S. pombe*.

Finally, we wanted to test whether gene function can also be used to predict protein abundance. We used the same linear equation and random walk as above, but performed the random walk with protein abundance values (Lawless *et al*. 2016) in place of RPKM values. As the random walk progressed over 10^5^ iterations, the correlation increased up to a maximum of 0.62 (Figure S13), a correlation that is even higher than the one generated using RPKM values in the random walk. The random walk was initiated 10 independent times with randomly-chosen β coefficients, generating the same correlation and β coefficients each time. For example, Figure S14 shows a linear relationship between the β coefficients of the 1^st^ and 5^th^ repeats. The β coefficients obtained with protein data versus RPKM data were somewhat consistent (compare Figure S9 with Figure S14). In both analyses, the β coefficient for “generation of precursor metabolites and energy” was among the highest and that of “cellular respiration” was among the lowest.

## DISCUSSION

RNA and protein levels increase or decrease upon changing cellular conditions, giving rise to the concept of differential expression. This concept is important in understanding tissue-and condition-specific gene expression and is used to determine which gene functions are important in a given environment. In contrast, we focus here not on changes in expression, but on absolute steady-state abundances of mRNA and protein. According to cost-benefit analysis, the abundance of each gene product should be controlled (Wagner 2005; Dekel and Alon 2005; Lang *et al*. 2009). The costs are two-fold: energy consumed during transcription and translation as well as mass that is added to the already-packed volume of the cell (Dill *et al*. 2011). The benefit is to perform a necessary cellular function. There is a balance between cost and benefit, resulting in a steady-state set point that provides maximal fitness for the cell and organism. Indeed, we have found here that, at least in one condition, there is an expression set point for each gene. This leads to the question of what factors determine the set point for a gene. We hypothesized that the function of the protein product would be an important determinant. We could test this by employing the gene ontology (GO) framework which systematically describes gene function. Specifically, we predicted that genes sharing a GO category would exhibit similar expression. The GO framework divides function into three domains: molecular function, biological process, and cellular component. First, proteins with the same molecular function need to be maintained at the same cellular concentration because they will have similar biochemical properties (e.g., K_m_, k_cat_) and work on substrates of similar concentrations. Supporting this idea, when bacteria are grown long-term in different levels of the substrate, lactose, the cells evolve to express a proportional level of the LacZ enzyme (Dekel and Alon 2005). Second, proteins that participate in the same biological process should be kept at the same concentration because they are components of a pathway with similar flux at each step. Proteins within the same cellular component should be of related abundances in order to achieve similar protein concentrations in a defined physical volume.

Several analyses presented here support the conclusion that gene function is an important factor in determining gene expression. First, we found that mRNA and protein levels are correlated between *S. cerevisiae* genes and their orthologous genes in *S. pombe*, showing that the expression level of functionally-related genes has been conserved over millions of years. This finding is consistent with cross-species correlations between other organisms (Schrimpf *et al*. 2009; Laurent *et al*. 2010; Khan *et al*. 2013). Second, we found that gene expression within several GO categories is significantly higher or lower than seen in the entire genome. Interestingly, genes involved in the Central Dogma follow a pattern. DNA-related genes are expressed the least, transcription-related genes are in the middle, and translation-related genes are expressed the most. As discussed above, this finding fits in with the amplification of biomolecules that occurs in the Central Dogma. Third, we were able to use GO terms alone in calculating gene expression levels. A linear model was created using *S. cerevisiae* GO and gene expression information, but then it was successfully used to predict *S. pombe* gene expression. Fourth, while we primarily relied on the plethora of high-quality RNA-seq data, we also performed analysis with protein data, obtaining similar results. This last point is critical since we assume that gene function is most closely associated with the abundance of proteins, the factors that directly perform the cellular functions. Consistently, protein levels are under greater evolutionary constraints than mRNA levels (Khan *et al*. 2013), likely because fitness relies more on the optimal protein level. However, protein abundance data is not always as accurate as mRNA abundance data and does not cover much of the genome (Vogel and Marcotte 2012; Liu *et al*. 2016). With future advances in measuring protein abundance, this study can be repeated with high-quality genome-wide protein data.

Our work here has shown that gene function (as defined by biological process, molecular function and cellular component) is a strong determinant of gene expression level. This has implications for how gene expression has evolved within the biochemical constraints of a cell. The constraints (e.g., organelle volume, substrate concentration, optimal flux through each pathway, and the energy requirements of transcription and translation) governing the expression of each protein can be estimated by the protein’s associated GO categories. However, GO categories alone might not accurately capture these constraints. The categories are proxies for other biochemical features of the protein. In this case, it might be important to determine whether more specific features, like K_m_ or cellular substrate concentration, are important in driving gene expression levels.

For gene function, we used the GO Slim annotations at the Saccharomyces Genome Database (SGD) (Cherry *et al*. 2012). While these annotations were useful for the initial study of gene function and gene expression, it would be useful to carry out future studies with the entire set of GO terms (Ashburner *et al*. 2000; Boyle *et al*. 2004). This will be especially useful for predicting gene expression as there would be additional information describing gene function. For example, instead of a gene simply associated with “transcription factor binding,” the gene may be labeled as being part of “the core TFIIH complex when it is part of the general transcription factor TFIIH,” a term that could be a better predictor of gene expression. In addition, as genes are further characterized, they will receive additional GO annotations that will improve the accuracy of prediction. As we observed here, genes associated with a larger number of GO categories could be more accurately predicted.

This analysis was performed primarily with expression data obtained from a particular strain of the yeast *S. cerevisiae*, grown in rich media. It was important to study expression in one condition to examine levels that are maintained at steady-state. However, our results may be biased by condition-specific effects. For example, a large set of genes is subject to glucose-repression under rich media conditions (Kayikci and Nielsen 2015) and thus would be labeled as poorly expressed simply because transcription was turned off. When these genes are relieved of glucose repression, they may be highly expressed. To deal with this issue, one could perform this analysis using the highest observed abundance for each gene. Practically, this could be done in *S. cerevisiae*, since genome-wide expression has been monitored across hundreds of conditions. Thus, the abundance value obtained for each gene would be the maximum and represent the level when the gene is “turned on.” This way, the expression level of all genes can be fairly compared. This could even be performed in other species, such as humans, that have a large number of both RNA-seq studies and GO annotations.

In predicting gene expression levels, we fit GO category information into a linear equation and optimized the coefficients with a random walk. We achieved a decent correlation (r=0.44) between prediction and observed, especially considering that the predictions were on a continuous scale and that we predicted the expression of 6,717 genes. However, other machine learning approaches may be more effective at estimating expression levels. These approaches include neural net, decision tree, naïve Bayes, and alternative mathematical models. The relationship between gene function and expression level is likely complex and further work is needed to determine the type of model that best takes into account all of the evolutionary forces that dictate gene expression levels.

## ACKNOWLEDGMENTS

This work was supported by funds from Rowan University and NIH 1R15GM113187 to M.J.H.

## LITERATURE CITED

Adhikari H., Cullen P. J., 2014 Metabolic Respiration Induces AMPK-and Ire1p-Dependent Activation of the p38-Type HOG MAPK Pathway. PLoS Genet 10: e1004734.

Anders S., Pyl P. T., Huber W., 2014 HTSeq A Python framework to work with high-throughput sequencing data.

Ashburner M., Ball C. A., Blake J. A., Botstein D., Butler H., Cherry J. M., Davis A. P., Dolinski K., Dwight S. S., Eppig J. T., Harris M. A., Hill D. P., Issel-Tarver L., Kasarskis A., Lewis S., Matese J. C., Richardson J. E., Ringwald M., Rubin G. M., Sherlock G., 2000 Gene ontology: tool for the unification of biology. The Gene Ontology Consortium. Nat Genet 25: 25–29.

Baker L. A., Ueberheide B. M., Dewell S., Chait B. T., Zheng D., Allis C. D., 2013 The yeast Snt2 protein coordinates the transcriptional response to hydrogen peroxide-mediated oxidative stress. Mol. Cell. Biol. 33: 3735–3748.

Bendjilali N., MacLeon S., Kalra G., Willis S. D., Hossian A. K. M. N., Avery E., Wojtowicz O., Hickman M. J., 2017 Time-Course Analysis of Gene Expression During the Saccharomyces cerevisiae Hypoxic Response. G3 (Bethesda) 7: 221–231.

Benjamini Y., Hochberg Y., 1995 Controlling the False Discovery Rate: A Practical and Powerful Approach to Multiple Testing on JSTOR. Journal of the Royal Statistical Society Series B ….

Blankenberg D., Gordon A., Kuster Von G., Coraor N., Taylor J., Nekrutenko A., Team G., 2010 Manipulation of FASTQ data with Galaxy. ….

Boyle E. I., Weng S., Gollub J., Jin H., Botstein D., Cherry J. M., Sherlock G., 2004 GO::TermFinder--open source software for accessing Gene Ontology information and finding significantly enriched Gene Ontology terms associated with a list of genes. Bioinformatics 20: 3710–3715.

Cherry J. M., Hong E. L., Amundsen C., Balakrishnan R., Binkley G., Chan E. T., Christie K. R., Costanzo M. C., Dwight S. S., Engel S. R., Fisk D. G., Hirschman J. E., Hitz B. C., Karra K., Krieger C. J., Miyasato S. R., Nash R. S., Park J., Skrzypek M. S., Simison M., Weng S., Wong E. D., 2012 Saccharomyces Genome Database: the genomics resource of budding yeast. Nucleic Acids Research 40: D700–5.

Conesa A., Madrigal P., Tarazona S., Gomez-Cabrero D., Cervera A., McPherson A., Szcześniak M. W., Gaffney D. J., Elo L. L., Zhang X., Mortazavi A., 2016 A survey of best practices for RNA-seq data analysis. Genome Biol 17: 13.

Csárdi G., Franks A., Choi D. S., Airoldi E. M., Drummond D. A., 2015 Accounting for experimental noise reveals that mRNA levels, amplified by post-transcriptional processes, largely determine steady-state protein levels in yeast. PLoS Genet 11: e1005206.

de Groot P. W. J., Brandt B. W., Horiuchi H., Ram A. F. J., de Koster C. G., Klis F. M., 2009 Comprehensive genomic analysis of cell wall genes in Aspergillus nidulans. Fungal Genetics and Biology 46: S72–S81.

Dekel E., Alon U., 2005 Optimality and evolutionary tuning of the expression level of a protein. Nature 436: 588–592.

Dill K. A., Ghosh K., Schmit J. D., 2011 Physical limits of cells and proteomes. Proc. Natl. Acad. Sci. U.S.A. 108: 17876–17882.

Dolinski K., Botstein D., 2007 Orthology and Functional Conservation in Eukaryotes. Annu. Rev. Genet. 41: 465–507.

Drawid A., Jansen R., Gerstein M., 2000 Genome-wide analysis relating expression level with protein subcellular localization. Trends Genet. 16: 426–430.

Engel S. R., Dietrich F. S., Fisk D. G., Binkley G., Balakrishnan R., Costanzo M. C., Dwight S. S., Hitz B. C., Karra K., Nash R. S., Weng S., Wong E. D., Lloyd P., Skrzypek M. S., Miyasato S. R., Simison M., Cherry J. M., 2014 The reference genome sequence of Saccharomyces cerevisiae: then and now. G3 (Bethesda) 4: 389–398.

Feijó Delgado F., Cermak N., Hecht V. C., Son S., Li Y., Knudsen S. M., Olcum S., Higgins J. M., Chen J., Grover W. H., Manalis S. R., 2013 Intracellular water exchange for measuring the dry mass, water mass and changes in chemical composition of living cells. PLoS ONE 8: e67590.

Fox M. J., Gao H., Smith-Kinnaman W. R., Liu Y., Mosley A. L., 2015 The Exosome Component Rrp6 Is Required for RNA Polymerase II Termination at Specific Targets of the Nrd1-Nab3 Pathway (J Corden, Ed.). PLoS Genet 11: e1004999.

Ghaemmaghami S., Huh W.-K., Bower K., Howson R. W., Belle A., Dephoure N., O’Shea E. K., Weissman J. S., 2003 Global analysis of protein expression in yeast. Nature 425: 737–741.

Jansen R., Gerstein M., 2000 Analysis of the yeast transcriptome with structural and functional categories: characterizing highly expressed proteins. Nucleic Acids Research 28: 1481–1488.

Jorgensen P., Edgington N. P., Schneider B. L., Rupes I., Tyers M., Futcher B., 2007 The size of the nucleus increases as yeast cells grow. Mol. Biol. Cell 18: 3523–3532.

Kayikci Ö., Nielsen J., 2015 Glucose repression in Saccharomyces cerevisiae. FEMS Yeast Research 15.

Khan Z., Ford M. J., Cusanovich D. A., Mitrano A., Pritchard J. K., Gilad Y., 2013 Primate transcript and protein expression levels evolve under compensatory selection pressures. Science 342: 1100–1104.

Kim D., Pertea G., Trapnell C., Pimentel H., Kelley R., Salzberg S. L., 2013 TopHat2: accurate alignment of transcriptomes in the presence of insertions, deletions and gene fusions. Genome Biol 14: R36.

Lang G. I., Murray A. W., Botstein D., 2009 The cost of gene expression underlies a fitness trade-off in yeast. Proc. Natl. Acad. Sci. U.S.A. 106: 5755–5760.

Laurent J. M., Vogel C., Kwon T., Craig S. A., Boutz D. R., Huse H. K., Nozue K., Walia H., Whiteley M., Ronald P. C., Marcotte E. M., 2010 Protein abundances are more conserved than mRNA abundances across diverse taxa. Proteomics 10: 4209–4212.

Lawless C., Holman S. W., Brownridge P., Lanthaler K., Harman V. M., Watkins R., Hammond D. E., Miller R. L., Sims P. F. G., Grant C. M., Eyers C. E., Beynon R. J., Hubbard S. J., 2016 Direct and Absolute Quantification of over 1800 Yeast Proteins via Selected Reaction Monitoring. Mol. Cell Proteomics 15: 1309–1322.

Leinonen R., Sugawara H., Shumway M., International Nucleotide Sequence Database Collaboration, 2011 The sequence read archive. Nucleic Acids Research 39: D19–21.

Li J. J., Chew G.-L., Biggin M. D., 2017 Quantitating Translational Control: mRNA Abundance-Dependent and Independent Contributions. bioRxiv: 116913.

Liu Y., Beyer A., Aebersold R., 2016 On the Dependency of Cellular Protein Levels on mRNA Abundance. Cell 165: 535–550.

Maier T., Güell M., Serrano L., 2009 Correlation of mRNA and protein in complex biological samples. FEBS Letters 583: 3966–3973.

Martín G. M., King D. A., Green E. M., Garcia-Nieto P. E., Alexander R., Collins S. R., Krogan N. J., Gozani O. P., Morrison A. J., 2014 Set5 and Set1 cooperate to repress gene expression at telomeres and retrotransposons. Epigenetics 9: 513–522.

McDowall M. D., Harris M. A., Lock A., Rutherford K., Staines D. M., Bähler J., Kersey P. J., Oliver S. G., Wood V., 2015 PomBase 2015: updates to the fission yeast database. Nucleic Acids Research 43: D656–61.

Mortazavi A., Williams B. A., McCue K., Schaeffer L., Wold B., 2008 Mapping and quantifying mammalian transcriptomes by RNA-Seq. Nat Meth 5: 621–628.

Mukherjee K., Gardin J., Futcher B., Leatherwood J., 2016 Relative contributions of the structural and catalytic roles of Rrp6 in exosomal degradation of individual mRNAs. RNA 22: 1311–1319.

Nagalakshmi U., Wang Z., Waern K., Shou C., Raha D., Gerstein M., Snyder M., 2008 The transcriptional landscape of the yeast genome defined by RNA sequencing. Science 320: 1344–1349.

Ozcan S., Johnston M., 1999 Function and regulation of yeast hexose transporters. Microbiol. Mol. Biol. Rev. 63: 554–569.

Risso D., Schwartz K., Sherlock G., Dudoit S., 2011 GC-content normalization for RNA-Seq data. BMC Bioinformatics 12: 480.

Schrimpf S. P., Weiss M., Reiter L., Ahrens C. H., Jovanovic M., Malmström J., Brunner E., Mohanty S., Lercher M. J., Hunziker P. E., Aebersold R., Mering von C., Hengartner M. O., 2009 Comparative functional analysis of the Caenorhabditis elegans and Drosophila melanogaster proteomes. Plos Biol 7: e48.

Shah M., Su D., Scheliga J. S., Pluskal T., Boronat S., Motamedchaboki K., Campos A. R., Qi F., Hidalgo E., Yanagida M., Wolf D. A., 2016 A Transcript-Specific eIF3 Complex Mediates Global Translational Control of Energy Metabolism. Cell Rep 16: 1891–1902.

Sipiczki M., 2000 Where does fission yeast sit on the tree of life? Genome Biol 1: REVIEWS1011.

Spellman P. T., Sherlock G., Zhang M. Q., Iyer V. R., Anders K., Eisen M. B., Brown P. O., Botstein D., Futcher B., 1998 Comprehensive identification of cell cycle-regulated genes of the yeast Saccharomyces cerevisiae by microarray hybridization. Mol. Biol. Cell 9: 3273–3297.

Team R. C., 2015 R: A Language and Environment for Statistical Computing. URL: http://www.R-project.org, Vienna.

Vaquerizas J. M., Kummerfeld S. K., Teichmann S. A., Luscombe N. M., 2009 A census of human transcription factors: function, expression and evolution. Nature Reviews Genetics 10: 252–263.

Velculescu V. E., Zhang L., Zhou W., Vogelstein J., Basrai M. A., Bassett D. E., Hieter P., Vogelstein B., Kinzler K. W., 1997 Characterization of the yeast transcriptome. Cell 88: 243–251.

Vogel C., 2013 Evolution. Protein expression under pressure. Science 342: 1052–1053.

Vogel C., Marcotte E. M., 2012 Insights into the regulation of protein abundance from proteomic and transcriptomic analyses. Nature Reviews Genetics 13: 227–232.

Wagner A., 2005 Energy Constraints on the Evolution of Gene Expression. Mol. Biol. Evol. 22: 1365–1374.

Wang M., Herrmann C. J., Simonovic M., Szklarczyk D., Mering von C., 2015 Version 4.0 of PaxDb: Protein abundance data, integrated across model organisms, tissues, and cell-lines. Proteomics 15: 3163–3168.

Wittkopp P. J., 2014 Evolution of Gene Expression. In:Losos JB, Baum DA, Futuyma DJ, Hoekstra HE, Lenski RE, Moore AJ, Peichel CL, Schluter D, Whitlock MJ (Eds.), The Princeton Guide to Evolution, Princeton University Press, Princeton, pp. 413–419.

Wood V., Harris M. A., McDowall M. D., Rutherford K., Vaughan B. W., Staines D. M., Aslett M., Lock A., Bähler J., Kersey P. J., Oliver S. G., 2012 PomBase: a comprehensive online resource for fission yeast. Nucleic Acids Research 40: D695–9.

